# Achilles tendon stiffness minimizes the energy cost in simulations of walking in older but not in young adults

**DOI:** 10.1101/2020.02.10.941591

**Authors:** Tijs Delabastita, Friedl De Groote, Benedicte Vanwanseele

## Abstract

Both Achilles tendon stiffness and walking patterns influence the energy cost of walking, but their relative contributions remain unclear. These independent contributions can only be investigated using simulations. We created models for 16 young (24±2 years) and 15 older (75±4 years) subjects, with individualized (using optimal parameter estimations) and generic triceps surae muscle-tendon parameters. We varied Achilles tendon stiffness and calculated the energy cost of walking. Both in young and older adults, Achilles tendon stiffness independently contributed to the energy cost of walking. However, overall, a 25% increase in Achilles tendon stiffness increased the triceps surae and whole-body energy cost of walking with approximately 7% and 1.5%, respectively. Therefore, the influence of Achilles tendon stiffness is rather limited. Walking patterns also independently contributed to the energy cost of walking because the plantarflexor (including, but not limited to the triceps surae) energy cost of walking was lower in older than in young adults. Hence, training interventions should probably rather target specific walking patterns than Achilles tendon stiffness to decrease the energy cost of walking. However, based on the results of previous experimental studies, we expected that the calculated hip extensor and whole-body energy cost of walking would be higher in older than in young adults. This was not confirmed in our results. Future research might therefore assess the contribution of the walking pattern to the energy cost of walking by individualizing maximal isometric muscle force and by using three-dimensional models of muscle contraction.

**Summary statement:** Achilles tendon stiffness and walking patterns independently contribute to the energy cost in simulations of walking in young and older adults. The influence of Achilles tendon stiffness is rather small.

## Introduction

The energy cost of walking is higher in older than in young adults (Malatesta et al., 2003) but the underlying mechanisms remain unclear (VanSwearingen and Studenski, 2014). In general, the energy cost of walking is largely determined by the skeletal muscles’ energy cost (Workman and Armstrong, 1986). The skeletal muscles’ energy cost is determined by muscle forces, activations, and contractile conditions, i.e. operating length and operating velocity (Umberger et al., 2003). Specific walking patterns impose joint torque demands that determine muscle forces during walking. Both specific walking patterns and the skeletal muscles’ in-series tendon stiffness influence muscle activations and contractile conditions. The influence of tendon stiffness on the muscle activations and contractile conditions is larger for muscles with longer and elastic energy-storing tendons, e.g. the Achilles tendon. Interestingly, both Achilles tendon stiffness (Delabastita et al., 2020) and the walking pattern (Boyer et al., 2017) are found to differ in young and older adults. Although their influence to the energy cost of walking is related (Delabastita et al., 2020), most studies only evaluated either the influence of Achilles tendon stiffness or the influence of specific walking patterns on the energy cost of walking. Therefore, the relative contributions of alterations in Achilles tendon stiffness versus alterations in walking patterns are not well understood.

Achilles tendon stiffness could influence the energy cost of walking through its influence on triceps surae muscle contractile conditions, which in turn determine triceps surae muscle activations required for a given force output (Lichtwark and Wilson, 2007; Lichtwark and Wilson, 2008) through the force-length (Gordon et al., 1966) and force-velocity (Hill, 1938) relationships of muscles. In young adults, it has been suggested that altering Achilles tendon stiffness would increase the energy cost of walking based on experimentally measured triceps surae muscle contractile conditions (Fukunaga et al., 2001). These contractile conditions, i.e. nearly isometric and near optimal fiber length, were found to minimize muscle energy cost of force production (Gordon et al., 1966; Ryschon et al., 1997). In older adults, it has been suggested that increasing Achilles tendon stiffness would decrease the energy cost of walking based on the observed correlation between Achilles tendon stiffness and maximal walking capacity measured by the six-minute walk distance (Stenroth et al., 2015).

Age-related differences in walking patterns associated with decreased triceps surae muscle strength might explain age-related differences in energy cost of walking. Older adults walk with a more flexed walking pattern than young adults (Boyer et al., 2017). This walking pattern requires lower plantarflexor moments but higher hip extensor moments compared to young adults’ walking pattern (Boyer et al., 2017). This altered walking pattern might increase the energy cost of walking (Wert et al., 2010), possibly due to higher energy dissipation when redirecting the body center of mass during step-to-step transitions (Collins and Kuo, 2010). As an altered walking pattern is related to lower maximal plantarflexor force (Judge et al., 1996), an age-related decrease in maximal plantarflexor force might partly drive the selection of an energetically less efficient walking pattern (Franz, 2016).

Since age-related changes in Achilles tendon stiffness and the walking pattern often occur simultaneously, it is very hard to design experimental studies that evaluate the causal contributions of alterations in Achilles tendon stiffness or the walking pattern to the energy cost of walking. However, simulations allow evaluating the isolated effect of either Achilles tendon stiffness or a specific walking pattern on the energy cost of walking. Inverse dynamic simulations use measured walking patterns and a musculoskeletal model as an input and calculate the underlying muscle fiber lengths, muscle activations and muscle forces. To do so, the muscle redundancy problem, the problem that arises from the fact that many more muscles than degrees of freedom exist, is solved optimizing a movement related cost function. Often, the sum of muscle activations squared is optimized (Crowninshield and Brand, 1981). Calculated fiber lengths, activations and forces can be used as inputs to metabolic energy models to calculate the whole-body and triceps surae energy cost of walking (Bhargava et al., 2004; Uchida et al., 2016; Umberger, 2010; Umberger et al., 2003). In such simulations, both walking patterns and Achilles tendon stiffness can be varied independently to evaluate their effect on the triceps surae muscle and whole-body energy cost of walking.

The primary aim of this study is to evaluate relative contributions of Achilles tendon stiffness and the specific walking pattern to the energy cost of walking in young and older adults. We applied an inverse dynamic approach using measured walking patterns and different musculoskeletal models as inputs. To evaluate the contribution of Achilles tendon stiffness to the energy cost of walking, we used musculoskeletal models with individualized triceps surae muscle-tendon parameters and varied Achilles tendon stiffness. We individualized triceps surae muscle-tendon parameters because calculated energy cost of walking is sensitive to changes in triceps surae muscle-tendon parameters (Delabastita et al., 2019). Hence, besides the influence of varying Achilles tendon stiffness, different triceps surae muscle-tendon parameters and different walking patterns might explain differences in the energy cost of walking. To evaluate the contribution of the walking pattern to the energy cost of walking, we used musculoskeletal models with scaled generic triceps surae muscle-tendon parameters. Hence, only differences in walking patterns would explain differences in calculated energy cost of walking. First, we expected that Achilles tendon stiffness would independently contribute to the energy cost of walking. We expected that decreasing and increasing Achilles tendon stiffness would increase the energy cost of walking in young adults. We also expected that decreasing and increasing Achilles tendon stiffness would, respectively, increase and decrease the energy cost of walking in older adults. Second, we expected that specific walking patterns in young and older adults would independently contribute to the energy cost of walking. Since plantarflexor moments are lower and hip extensor moments are higher in older than in young adults, we expected that the plantarflexor energy cost would be lower and hip extensor energy cost would be higher in older than in young adults. Furthermore, since the measured whole-body energy cost of walking is higher in older than in young adults (Malatesta et al., 2003), we expected that the whole-body energy cost of walking calculated using inverse simulations would also be higher in older than in young adults.

Different methods to calculate muscle and whole-body energy cost of walking based on muscle fiber lengths, muscle activations, and muscle forces have been proposed (Bhargava et al., 2004; Uchida et al., 2016; Umberger, 2010; Umberger et al., 2003), but it is unclear which method calculates the energy cost of walking most accurately since experimental validation is lacking. Such experimental validation, comparing measured and calculated energy cost of walking using different methods, can only be performed at the level of the whole-body energy cost of walking that can be measured using indirect calorimetry (Wert et al., 2015) because it is infeasible to measure the triceps surae energy cost of walking (Marsh and Ellerby, 2006). A previous validation study showed that these methods better predict within-subject differences than between-subject differences, however only two-dimensional musculoskeletal models with a very limited amount of muscle actuators were used (Koelewijn et al., 2019).

Therefore, the secondary aim of this study is to evaluate which metabolic energy model most accurately predicts within-subjects changes in energy cost of walking due to changes in walking speed using three-dimensional musculoskeletal models. We compared different models to select the most accurate model to evaluate relative contributions of specific walking patterns and Achilles tendon stiffness to the energy cost of walking in young and older adults.

## Methods

To evaluate the relative contribution of walking patterns and Achilles tendon stiffness to the energy cost of walking in young and older adults, we used inverse dynamic simulations with measured walking patterns and different musculoskeletal models, i.e. with individualized and generic triceps surae muscle-tendon parameters, to calculate the energy cost using different simulation-based methods. As inputs to inverse dynamics simulations, we quantified joint moments, muscle-tendon lengths and muscle-tendon moment arms using marker data and ground reaction forces. To individualize triceps surae muscle-tendon parameters using optimal parameter estimations, we additionally quantified muscle activations using EMG data, gastrocnemius medialis fiber length, and gastrocnemius medialis pennation angle using ultrasound images during walking. To evaluate which method most accurately predicts within-subjects changes in energy cost of walking due to changes in walking speed, we quantified the energy cost of walking using indirect calorimetry.

### Experimental design

Sixteen healthy young (age 24±2 years, mass 64±12 kg, and length 170±9 cm) and eighteen healthy older adults (age 75±4 years, mass 73±12 kg, and length 165±7 cm) participated in this study. We included subjects above 68 years (older) and between 20 and 30 years old (young). These subjects did not participate in competitive sports or did not perform structured exercise more than twice a week over the last three months. We excluded three older subjects from further analysis. Two subjects were excluded due to insufficient ultrasound image quality and one subject due to technical problems using indirect calorimetry. The Medical Ethics Committee UZ/KU Leuven approved the study and subjects signed written consent prior to participation in accordance with the Declaration of Helsinki.

To evaluate within-subject changes in energy cost of walking across a broad range of walking speeds, we determined the subject’s comfortable walking speed and the subject’s mean speed during a six-minute walk test as previously described (Delabastita et al., 2019). Next, participants familiarized with walking on the instrumented treadmill (Motekforce Link, Amsterdam, The Netherlands) for at least 20 minutes. Afterwards, we prepared the subjects and we started actual treadmill testing. During treadmill walking, the subjects walked at 3 km/h (slow), 5 km/h (fast), their comfortable walking speed and their six-minute walk speed. We collected marker data, ground reaction forces, EMG data, and ultrasound data during three consecutive strides after five to eight minutes of walking at a specific walking speed when subjects attained a steady energy consumption. Afterwards, we proceeded to the next walking speed.

### Data collection

To quantify joint moments, muscle-tendon lengths and muscle-tendon moment arms, we collected marker data and ground reaction forces during walking. We captured the trajectories of 68 reflective markers using thirteen infrared cameras (Vicon, Oxford, UK) recording at 100 Hz. We used a full body plug in gait marker set extended with 4 upper limb clusters, 4 lower limb clusters and one pelvis cluster (Van Rossom et al., 2017). We collected ground reaction forces using two force plates built in the treadmill at 1000 Hz.

To quantify muscle activations, we collected EMG signals of the right gastrocnemius lateralis, soleus and tibialis anterior at 1000 Hz (ZeroWire EMG, Cometa, Milano, Italy). To quantify gastrocnemius medialis fiber length and pennation angle during walking, we collected ultrasound images of the left gastrocnemius medialis with a PC-based ultrasound system (Echoblaster 128, UAB Telemed, Vilnius, Lithuania). We used a flat shaped probe with an 8 MHz wave frequency and a 60 mm field of view sampling at 60 Hz. We carefully aligned the probe with the muscle fascicles and the deep aponeurosis of the gastrocnemius medialis (Bolsterlee et al., 2016). We synchronized ultrasound and motion capture data with an electrical signal during ultrasound recording.

To quantify the energy cost of walking using indirect calorimetry, we collected the subject’s oxygen consumption and carbon dioxide production during treadmill walking continuously (Oxycon Mobile Pro, Carefusion, Houten, The Netherlands). We instructed the subjects to maintain a normal dietary pattern and to avoid intense physical activity during 24 hours before the start of the measurement. The subjects did not eat or drink for at least two hours before treadmill testing started.

### Data processing

As inputs to inverse dynamics simulations, we quantified joint moments, muscle-tendon lengths and muscle-tendon moment arms during walking. We scaled an OpenSim gait2392-model using static standing marker data (Delp et al., 2007). Consequently, we used the scaled model and the treadmill walking marker data to compute joint kinematics with a Kalman smoothing algorithm (De Groote et al., 2008). We used the joint kinematics to calculate muscle-tendon lengths and moment arms with OpenSim’s muscle analysis tool. We also used the kinematics and the ground reaction forces to calculate joint moments during walking with OpenSim’s inverse dynamics tool.

To individualize triceps surae muscle-tendon parameters using optimal parameter estimations, we additionally quantified muscle activations, gastrocnemius medialis fiber length, and gastrocnemius medialis pennation angle. To quantify muscle activations during walking, we band-pass filtered (20-400 Hz), rectified and low-pass filtered (10 Hz) the EMG signals of the right gastrocnemius lateralis, soleus and tibialis anterior. To quantify gastrocnemius medialis fiber length and pennation angle during walking, we tracked three lines representing the superficial aponeurosis, the deep aponeurosis and the orientation of the gastrocnemius medialis muscle fascicles with a semi-automated algorithm (Farris, D.J., Lichtwark, 2016). We linearly extrapolated these lines and calculated gastrocnemius medialis fiber length as the distance between the two aponeuroses parallel to the muscle fascicles and the pennation angle as the angle between the muscle fascicles and the deep aponeurosis (Lichtwark et al., 2007).

To evaluate which method most accurately predicts within-subjects changes in energy cost of walking due to changes in walking speed, we quantified the energy cost of walking. Since calculated whole-body energy cost is the sum of all muscles’ energy cost, we quantified the net energy consumption by subtracting the average energy consumption during static standing (Ortega and Farley, 2007) from the average whole-body energy consumption during the last two minutes of treadmill walking (Oxycon Mobile Pro software, Carefusion, Houten, The Netherlands). To quantify the (net) measured energy cost of walking, we normalized the net energy consumption during walking to body weight and to walking speed (Malatesta et al., 2003).

#### Optimal parameter estimation

The subject’s scaled generic model, muscle-tendon lengths, muscle moment arms, joint moments, measured gastrocnemius medialis fiber length and pennation angle, and measured muscle activations were inputs to an recently developed optimal parameter estimation algorithm (Delabastita et al., 2019) (Fig. 1, full lines and textboxes). The algorithm estimated Achilles tendon stiffness, gastrocnemius medialis and lateralis optimal fiber length, gastrocnemius medialis and lateralis tendon slack length and gastrocnemius medialis pennation angle at optimal fiber length. It estimated these triceps surae muscle-tendon parameters by optimizing the fit between measured and calculated gastrocnemius medialis fiber length and pennation angle, and measured and calculated gastrocnemius lateralis, soleus, and tibialis anterior muscle activations. We estimated the parameters simultaneously at one stride of each walking speed to compute one set of estimated parameters that was optimal for all walking speeds in our dataset.

**Figure 1.**
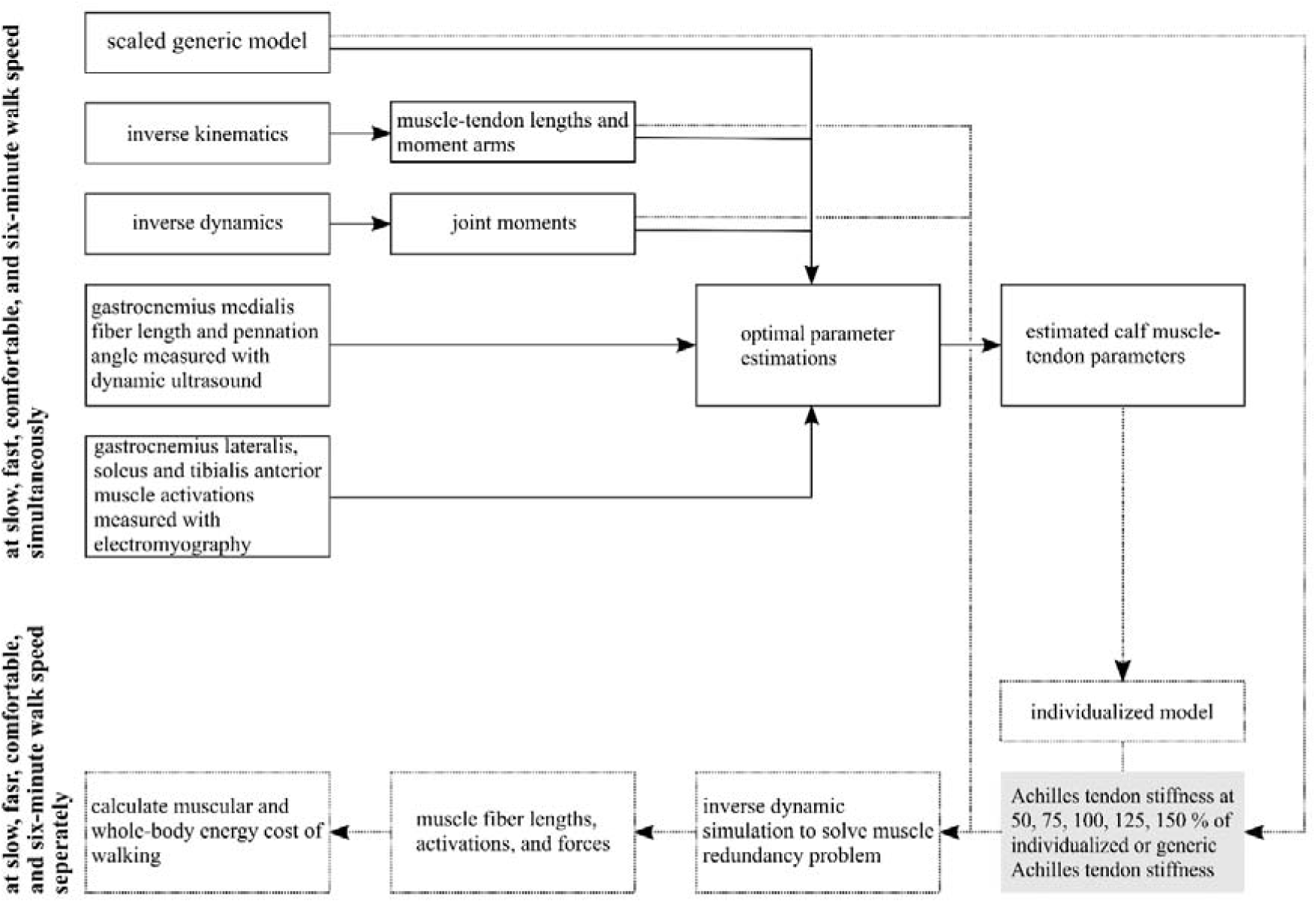
Workflow to calculate the metabolic energy cost of walking using model with different triceps surae muscle-tendon parameters (individualized with optimal parameter estimations or generic) and with different variations in Achilles tendon stiffness in young and older adults. We estimated the parameters gastrocnemius medialis optimal fiber length, tendon slack length and pennation angle at optimal fiber length, and gastrocnemius lateralis optimal fiber length and tendon slack length, and Achilles tendon stiffness.

#### *The relative contribution of Achilles tendon stiffness and specific walking patterns to the energy cost of walking using models with individualized and generic* triceps surae *muscle-tendon parameters*

To evaluate the contribution of Achilles tendon stiffness to the energy cost of walking, we used musculoskeletal models with individualized triceps surae muscle-tendon parameters and varied Achilles tendon stiffness because calculated energy cost is sensitive to changes in triceps surae muscle-tendon parameters (Delabastita et al., 2019). We used the model with individualized triceps surae muscle-tendon parameters to create additional models by changing Achilles tendon stiffness to 50%, 75%, 125% and 150% of individualized Achilles tendon stiffness (Fig. 1, grey textbox). Using models with individualized triceps surae muscle-tendon parameters, both the effects of individualized triceps surae muscle-tendon parameters and specific walking patterns in young and older adults mediate the effect of varying Achilles tendon stiffness on muscle fiber lengths, activations, and forces.

To evaluate the contribution of specific walking patterns to the energy cost of walking, we used musculoskeletal models with scaled generic triceps surae muscle-tendon parameters (Fig. 1, grey textbox, 100% of generic Achilles tendon stiffness). Using models with generic triceps surae muscle-tendon parameters, only specific walking patterns in young and older adults would explain differences in muscle fiber lengths, activations, and forces. To be able to compare the influence of varying Achilles tendon stiffness in models with individualized and generic Achilles tendon stiffness, we set generic Achilles tendon stiffness to 26. Although previous studies used 35 (Zajac, 1989), we adapted it to 26 because this is the mean of estimated Achilles tendon stiffness across all subjects. This value better represents the subjects in our study sample, and enables better comparisons to the outcomes calculated with (variations to) individualized Achilles tendon stiffness.

Additionally, to evaluate if specific walking patterns and other triceps surae muscle-tendon parameters influence the contribution of Achilles tendon stiffness to the energy cost of walking, we varied Achilles tendon stiffness in a model with generic triceps surae muscle-tendon parameters. We created additional models by changing Achilles tendon stiffness to 50%, 75%, 125% and 150% of generic Achilles tendon stiffness (Fig. 1, grey textbox). If the differences between young and older adults in the effect of varying Achilles tendon stiffness on the energy cost of walking are equal when using models with generic instead of individualized triceps surae muscle-tendon parameters, we can attribute differences between young and older adults to differences in walking patterns. If differences between young and older adults in the effect of varying Achilles tendon stiffness on the energy cost of walking change when using models with generic instead of individualized triceps surae muscle-tendon parameters, we can attribute differences between young and older adults to differences in triceps surae muscle-tendon parameters.

As such, for every subject, we created ten scaled musculoskeletal models with varying (including individualized and generic) Achilles tendon stiffness and with individualized and generic gastrocnemius medialis and lateralis fiber length, gastrocnemius medialis and lateralis tendon slack length, gastrocnemius medialis pennation angle at optimal fiber length.

#### Solving the muscle redundancy problem

Consequently, we used individualized and scaled generic models with varying Achilles tendon stiffness, muscle-tendon lengths, muscle-tendon moment arms, joint moments during walking to solve the muscle redundancy problem to calculate muscle forces, fiber lengths and activations underlying treadmill walking at each walking speed (Fig. 1, dotted lines and textboxes). We applied an existing dynamic optimization approach for inverse dynamic simulations (De Groote et al., 2016).

#### Calculate the energy cost of walking

To calculate the energy cost of walking, we applied different methods using muscle forces, fiber lengths and activations as inputs (Bhargava et al., 2004; Uchida et al., 2016; Umberger, 2010; Umberger et al., 2003). These methods calculate the rate of muscular energy consumption as the sum of heat production due to activations and maintenance, heat production due to shortening and lengthening, and mechanical work production.

We took the time integral of each muscle’s metabolic energy consumption rate over the whole stride to quantify each muscle’s energy consumption during walking. We normalized each muscle’s energy consumption to body weight and distance covered (Malatesta et al., 2003) to quantify the muscle energy cost of walking.

To analyze the contribution of Achilles tendon stiffness to the energy cost of walking, we calculated the whole-body energy cost of walking as the sum of all muscles’ energy cost. We also calculated the triceps surae energy cost of walking. Additionally, we calculated the energy cost of all muscles or all plantarflexor muscles except the triceps surae to investigate if varying Achilles tendon stiffness would lead to compensations in other muscles. Moreover, we calculated the triceps surae heat production due to activations and maintenance, heat production due to shortening and lengthening, and mechanical work production (Umberger et al., 2003). We also calculated Achilles tendon positive work because mechanical tendon work reduces muscle fiber work during walking (Lichtwark et al., 2007).

To analyze the contribution of the walking pattern to the energy cost of walking, we used the whole-body energy cost of walking and additionally calculated the plantarflexor and hip extensor muscle energy cost of walking by summing all plantarflexor and hip extensor muscles’ energy cost of walking, respectively.

### Statistical analysis

To assess which method most accurately calculates the energy cost of walking, we computed the correlation between measured and calculated energy cost of walking across all four walking speeds for every method separately. We used musculoskeletal models with individualized triceps surae muscle-tendon parameters. We compared the correlations of the separate methods using a Kruskal-Wallis test to account for their non-normal distribution. We set the critical *α*-level at 0.05. We performed these analyses in SPSS (SPSS Statistics version 20, IBM, New York, USA).

To evaluate age-related differences in walking patterns, we compared joint angles, joint moments, comfortable and six-minute walk speed, and stride frequency at all walking speeds in young and older adults. To compare joint angles and joint moments in young and older adults, we used statistical non-parametric mapping that accounted for non-normally distributed variables (Pataky et al., 2015). We used a two-samples t-test to compute t-statistics and critical t-values over the whole stride (Pataky et al., 2015). At certain phases during the stride, the t-statistics exceed the threshold t-value, indicating a significant difference during these phases. We performed these analyses in MatLab 2015b (The Mathworks, Inc., Natick, USA). To compare comfortable and six-minute walk speed, stride frequency and measured energy cost of walking at all walking speeds in young and older adults, we used Mann-Whitney U tests that also accounted for their non-normal distribution. We performed these analyses in SPSS.

To compare estimated muscle-tendon parameters between young and older adults, we used Mann-Whitney U tests to account for their non-normal distribution. We also performed these analyses in SPSS.

To evaluate the contribution of Achilles tendon stiffness to the energy cost of walking, and to evaluate if specific walking patterns and other triceps surae muscle-tendon parameters influence the contribution of Achilles tendon stiffness to the energy cost of walking, we used generalized estimating equations to account for non-normal distributions. Energy cost (different outcomes described above) was included as the dependent variable; group was included as a between-subjects factor with two levels, and walking speed and variation in Achilles tendon stiffness were included as within-subjects factors with four and five levels, respectively. We applied Sidak-corrections for multiple comparisons. Hence, to evaluate the contribution of Achilles tendon stiffness to the energy cost of walking, we set the critical *α*-level for a significant interaction effect at 0.0085 (six comparisons). To evaluate if specific walking patterns and other triceps surae muscle-tendon parameters influence the contribution of Achilles tendon stiffness to the energy cost of walking, we set the critical *α*-level for a significant interaction effect at 0.0253 (two comparisons).

To evaluate the contribution of the walking pattern to the energy cost of walking, we also used generalized estimating equations to account for non-normal distributions. Energy cost (different outcomes described above) was included as the dependent variable; group was included as a between-subjects factor with two levels, and walking speed (but not variation in Achilles tendon stiffness) was included as a within-subjects factor with four levels. We also applied Bonferroni corrections for multiple pairwise comparisons and set the critical *α*-level for a significant interaction effect at 0.017 (three comparisons).

## Results

All metabolic energy models described the relationship between walking speed and whole-body energy cost of walking equally well (Fig. 2). The median correlations were 0.74 (Uchida et al., 2016), 0.75 (Bhargava et al., 2004), 0.78 (Umberger et al., 2003), and 0.83 (Umberger, 2010). Since the between-subjects range of the correlations was smallest for the model of (Umberger et al., 2003), we used this model in further analyses (Fig. 2).

**Figure 2.**
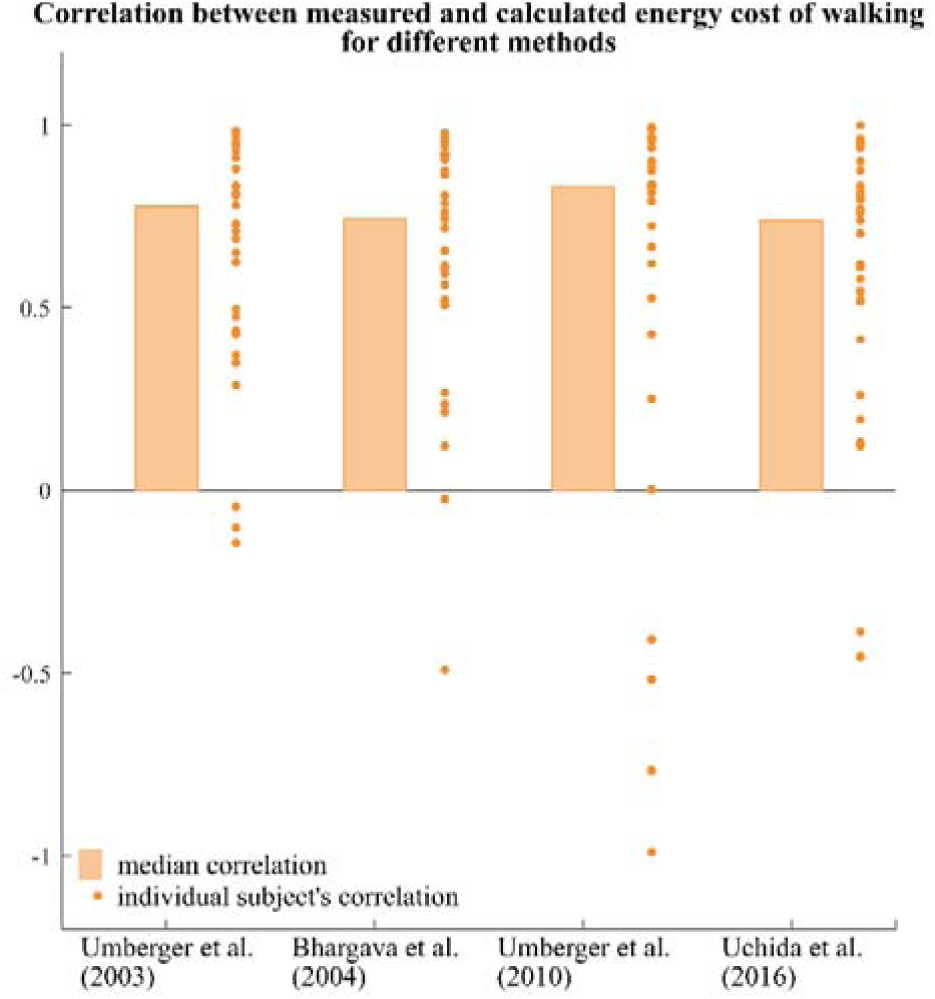
Correlation between measured and calculated energy cost of walking across four walking speeds for different methods to calculate the energy cost of walking in young and older adults together (n = 31).

Estimated triceps surae muscle optimal fiber length and walking patterns were different in young and older adults, which might influence the energy cost of walking. Gastrocnemius medialis and lateralis optimal fiber length was lower in older than in young adults (Table 1). Achilles tendon stiffness, gastrocnemius medialis tendon slack length and pennation angle at optimal fiber length, and gastrocnemius lateralis tendon slack length were equal in young and older adults (Table 1). Ankle plantarflexion angle during push-off was lower at all speeds and ankle plantarflexor moment during push-off was lower at all speeds, except at slow speed (Fig. S1). We only observed small differences in knee flexion angle and knee flexion moment in young and older adults that were not different across the different speeds (Fig. S2). Hip flexion during walking was higher in older than in young adults, and hip extensor moment during loading response and mid-stance were higher in older than in young adults (Fig. S3).

**Table 1.** Estimated triceps surae muscle-tendon parameters in young and older adults.

| Estimated parameter | Young adults | Older adults |
| --- | --- | --- |
| Normalized Achilles tendon stiffness [ ] | 26.3 ± 8.8 | 25.5 ± 5.4 |
| Gastrocnemius medialis optimal fiber length [cm] | 5.9 ± 0.1 | 4.9 ± 0.1* |
| Gastrocnemius medialis tendon slack length [cm] | 39.3 ± 3.1 | 38.4 ± 2.0 |
| Gastrocnemius medialis pennation angle at optimal fiber length [°] | 17.5 ± 2.8 | 17.1 ± 3.0 |
| Gastrocnemius lateralis optimal fiber length [cm] | **6.3 ± 0.1** | **5.2 ± 0.1\*** |
| Gastrocnemius lateralis tendon slack length [cm] | 38.3 ± 3.2 | 37.1 ± 2.1 |

Bold font and asterisks indicate significant differences between young and older adults.

Six-minute walk speed, stride frequency, and the energy cost of walking differed in young and older adults. Six-minute walk speed was lower in older than in young adults (Table 2). Stride frequency was higher at slow and fast speeds, but lower at six-minute walk speed in older compared to young adults (Table 2). The energy cost of walking was higher in young and older adults at slow and fast walking speeds.

**Table 2.** Walking speed, stride frequency, and measured energy cost of walking in young and older adults.

|  | Walking speed<br>[km/h] |  | Stride frequency<br>[strides/min] |  | Measured energy cost<br>[J/m/kg] |  |
| --- | --- | --- | --- | --- | --- | --- |
|  | YA | OA | YA | OA | YA | OA |
| Slow | 3.0 | 3.0 | **50.7 ± 2.9** | **55.6 ± 4.3\*** | **2.18 ± 0.33** | **2.58 ± 0.40\*** |
| Fast | 5.0 | 5.0 | **60.3 ± 3.0** | **63.9 ± 3.8\*** | **2.23 ± 0.37** | **2.54 ± 0.44\*** |
| Comfortable | 5.0 ± 0.6 | 4.6 ± 0.5 | 60.3 ± 4.1 | 61.6 ± 3.5 | 2.26 ± 0.32 | 2.45 ± 0.44 |
| Six-minute walk | **6.6 ± 0.6** | **5.4 ± 0.5 \*** | **68.2 ± 4.8** | **68.2 ± 3.1\*** | 3.09 ± 0.45 | 2.81 ± 0.60 |

Slow (3 km/h) and fast (5 km/h) speeds were matched in all subjects, while comfortable and six-minute walk speeds were individually determined. Bold font and asterisks indicate significant differences between young and older adults.

The effect of varying Achilles tendon stiffness on the energy cost of walking or the energy cost of walking itself were dependent on speed and age when using models with individualized and generic triceps surae muscle-tendon parameters (stiffness * group * age effect or group * age effects p < 0.001 for all parameters).

The influence of varying Achilles tendon stiffness on the whole-body (Fig. 3A) and triceps surae (Fig. 3B) energy cost of walking was different in young and older adults when using individualized triceps surae muscle-tendon parameters. In young adults, decreasing Achilles tendon stiffness decreased the whole-body energy cost of walking at fast and comfortable walking speeds and did not change or increased the whole-body energy cost of walking at slow and six-minute walk speed. Furthermore, in young adults, decreasing Achilles tendon stiffness decreased the whole-body energy cost of walking at fast, comfortable and six-minute walk speeds but and did not change the whole-body energy cost of walking at slow speed. In contrast, in young adults, increasing Achilles tendon stiffness increased or did not change whole-body and triceps surae energy cost of walking. Hence, in young adults, whole body or triceps surae energy cost of walking was minimal when calculated using a model with 50% or 75% of individualized Achilles tendon stiffness at fast and comfortable speed. In older adults, both decreasing and increasing Achilles tendon stiffness did not change or increased whole-body and triceps surae energy cost of walking. Hence, in older adults, whole body and triceps surae energy cost of walking were minimal when calculated using a model with individualized Achilles tendon stiffness at all speeds.

**Figure 3.**
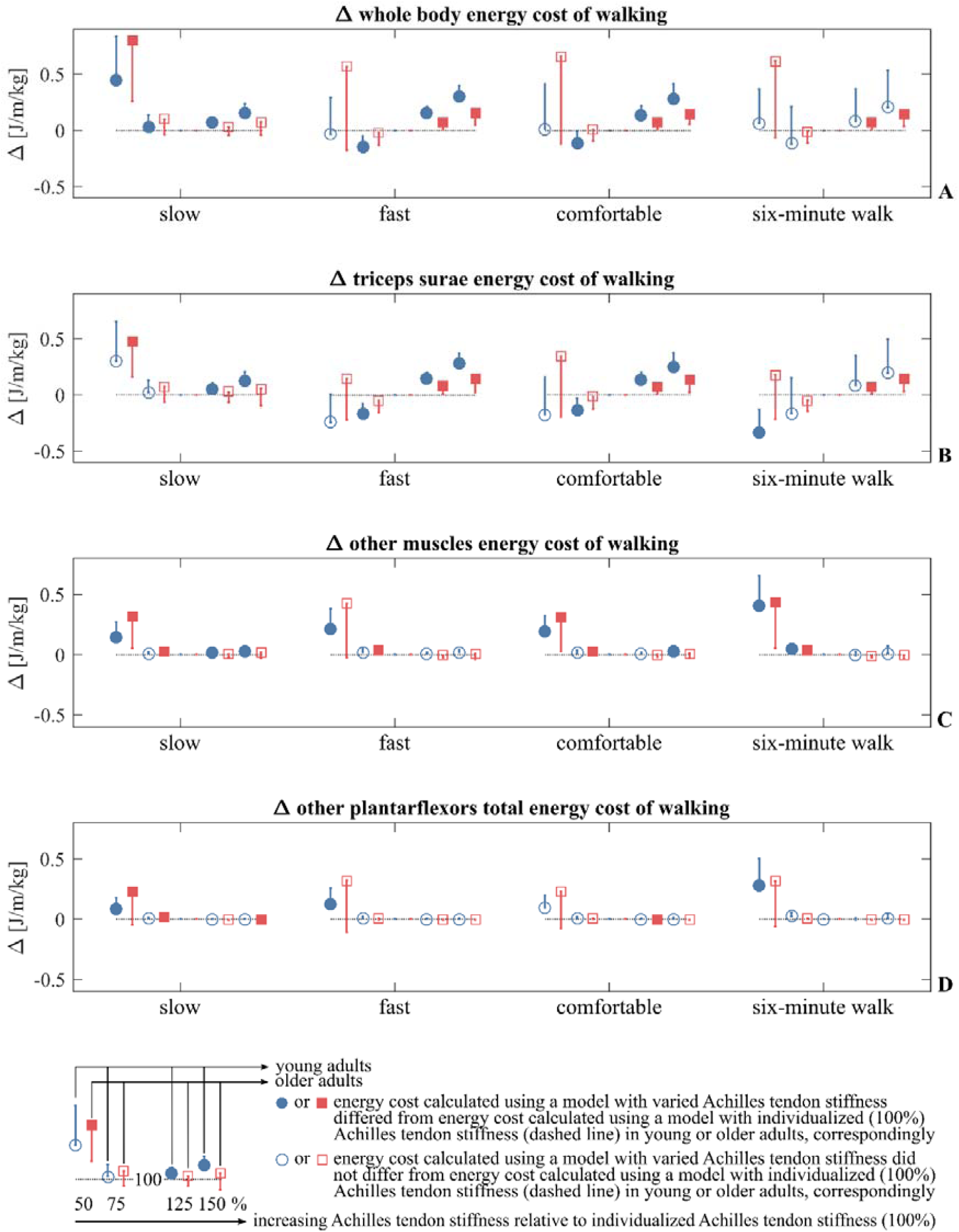
Influence of Achilles tendon stiffness on the energy cost of walking at different speeds calculated using models with individualized triceps surae muscle-tendon parameters and varying Achilles tendon stiffness in young and older adults. The differences in energy cost are expressed with respect to the model with individualized Achilles tendon stiffness. Error bars indicate standard deviations.

The other muscles’ (Fig. 3C) or other plantarflexor muscles’ (Fig. 3D) energy cost of walking slightly increased at 75% and clearly increased at 50% of individualized Achilles tendon stiffness in both young and older adults.

When calculated using models with generic triceps surae muscle-tendon parameters, the plantarflexor muscle energy cost of walking was lower in older than in young adults at most walking speeds (Fig. 4A) and the hip extensor energy cost of walking was equal in young and older adults at all speeds (Fig. 4B). The whole-body energy cost of walking was equal in young than in older adults at all speeds except six-minute walk speed. At six-minute walk speed, the whole-body energy cost of walking was lower in older than in young adults when calculated using models with generic triceps surae muscle-tendon parameters (Fig. 4C).

**Figure 4.**
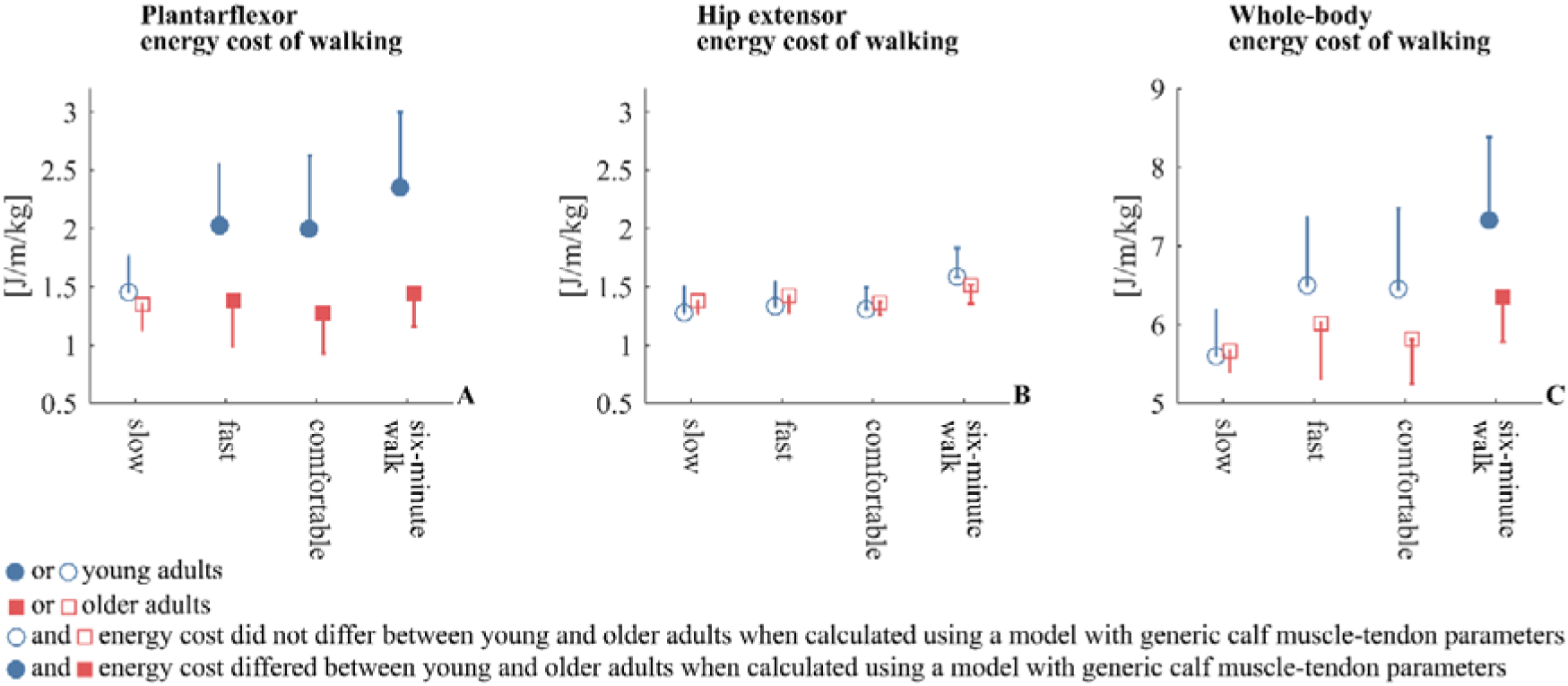
Influence of specific walking patterns on the energy cost of walking at different speeds calculated using models with generic triceps surae muscle-tendon parameters in young and older adults. Error bars indicate standard deviations.

With respect to the whole-body energy cost of walking, specific walking patterns in young and older adults mediate the effect of varying Achilles tendon stiffness. The differences between young and older adults in the effect of varying Achilles tendon stiffness on the whole-body energy cost of walking do not change when using models with generic instead of individualized triceps surae muscle-tendon parameters. For instance, at fast and comfortable walking speed, decreasing Achilles tendon stiffness decreases the whole-body energy cost of walking in young but not in older adults when calculated using both individualized (Fig. 3A) and generic (Fig. 5A) triceps surae muscle-tendon parameters.

**Figure 5.**
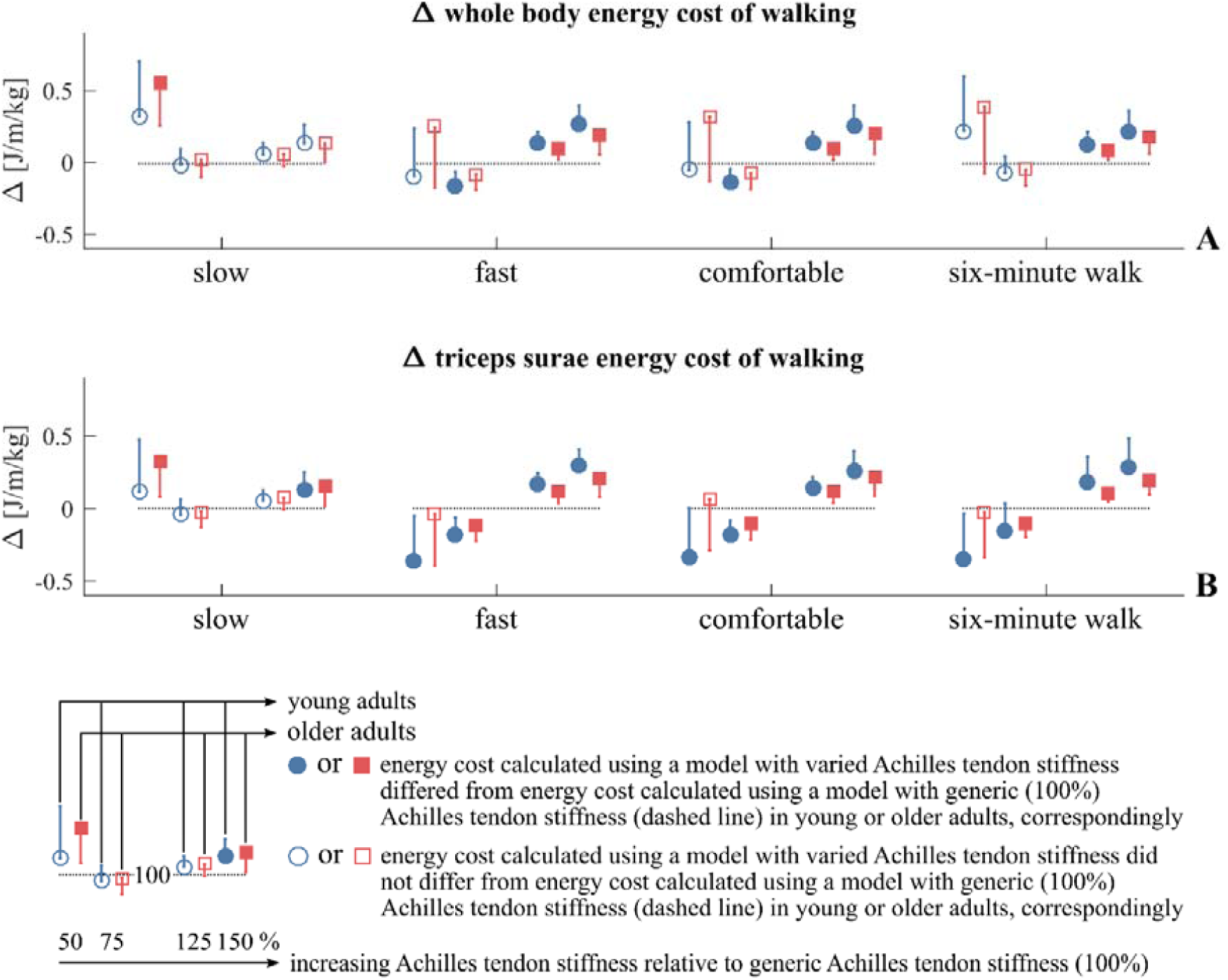
Influence of Achilles tendon stiffness on the energy cost of walking at different speeds calculated using models with scaled generic triceps surae muscle-tendon parameters and varying Achilles tendon stiffness in young and older adults. The differences in energy cost are expressed with respect to the model with individualized Achilles tendon stiffness. Error bars indicate standard deviations.

With respect to the triceps surae energy cost of walking, differences in optimal fiber length mediate the effect of varying Achilles tendon stiffness. The effect of varying Achilles tendon stiffness on the triceps surae energy cost of walking differs between young and older adults when using individualized but not when using generic triceps surae muscle-tendon parameters. For instance, at fast and comfortable walking speeds, decreasing Achilles tendon stiffness does not decrease the triceps surae energy cost of walking in older adults when calculated using individualized triceps surae muscle-tendon parameters (Fig. 3B), but it does decrease the triceps surae energy cost of walking in older adults when calculated using generic triceps surae muscle-tendon parameters (Fig. 5B). Similarly, decreasing Achilles tendon stiffness decreases the triceps surae energy cost of walking in young adults when calculated using both individualized (Fig. 3B) and generic (Fig. 5B) triceps surae muscle-tendon parameters.

## Discussion

The primary aim of this study is to evaluate the contribution of Achilles tendon stiffness and the walking patterns to the energy cost of walking in young and older adults. Previous studies only evaluated either the influence of specific walking patterns or the influence of Achilles tendon stiffness on the energy cost of walking. Furthermore, previous studies suggested that varying Achilles tendon stiffness would affect the triceps surae and whole-body energy cost of walking in young and older adults, but evidence for these suggestions was still lacking. Our study is the first to show that Achilles tendon stiffness and the walking pattern independently contribute to the triceps surae muscle and whole-body energy cost of walking, and that the interaction between specific walking patterns, Achilles tendon stiffness, and the energy cost of walking is different in young and older adults.

We could not confirm that increasing Achilles tendon stiffness decreases the energy cost of walking in older adults. In older adults, the calculated whole-body and triceps surae energy cost of walking were lowest with individualized Achilles tendon stiffness at all speeds. Therefore, we suggest that Achilles tendon stiffness minimizes the energy cost underlying older adults’ specific walking pattern. This is in contrast with a previous study suggesting that increasing Achilles tendon stiffness would decrease the energy cost of walking in older adults based on an independent correlation between Achilles tendon stiffness and maximal walking capacity measured using the six-minute walk distance (Stenroth et al., 2015). However, this study did not take into account the influence of age-related differences in walking patterns on triceps surae muscle contractile conditions (Stenroth et al., 2016).

We could not confirm that decreasing and increasing Achilles tendon stiffness both increase the energy cost of walking in young adults. In young adults, the whole-body and triceps surae energy cost of walking were lowest when calculated using a model with 75% of individualized Achilles tendon stiffness at fast and comfortable speeds. Therefore, it seems that decreasing Achilles tendon stiffness would decrease the energy cost underlying the specific walking patterns in young adults. This is in contrast with a study suggesting that both decreasing and increasing Achilles tendon stiffness would increase the energy cost of walking based on the experimental observation that triceps surae muscle fibers produce work at relatively constant length and low shortening velocities (Fukunaga et al., 2001). However, this study did not calculate the triceps surae or whole-body energy cost of walking.

Although Achilles tendon stiffness independently influences the energy cost underlying specific walking patterns in young and older adults, its influence is rather limited. A 25% increase in Achilles tendon stiffness, which is realistic because it is comparable to the standard deviation for estimated Achilles tendon stiffness (Table 1) and to increases in Achilles tendon stiffness following resistance training interventions (McCrum et al., 2018), increased the triceps surae and whole-body energy cost of walking with approximately 7% and 1.5%, respectively. Hence, training interventions should probably not target Achilles tendon stiffness to decrease the energy cost of walking.

Our results confirm that the walking pattern independently contributes to the energy cost of walking. As expected, the plantarflexor energy cost of walking was lower in older than in young adults, probably due to lower ankle plantarflexor moments at push-off in older than in young adults. In contrast to our expectations, the hip extensor energy cost of walking was equal in young and older adults. Maybe, we could not confirm that the hip extensor energy cost is higher in older than in young adults due to inaccuracies in our musculoskeletal models. Since muscle strength decreases with aging (Narici et al., 2008), individualizing maximal isometric muscle forces in our musculoskeletal models would increase the plantarflexor and hip extensor muscle activations and therefore the plantarflexor and hip extensor muscle energy cost of walking. Furthermore, as reported for the ankle plantarflexors (Rasske and Franz, 2018), it might be that the hip muscle moment arms during walking are smaller in older than in young adults due to smaller three-dimensional shape changes (muscle bulging). However, since the Hill-type muscle models assume constant muscle thickness, this is overlooked in our musculoskeletal models (Zajac, 1989). Hence, implementing a three-dimensional model of muscle contraction would increase the plantarflexor and hip extensor muscle forces during walking and therefore the plantarflexor and hip extensor energy cost of walking as well. If using models with individualized maximal isometric muscle forces and with three-dimensional models of muscle contraction would increase the plantarflexor and hip extensor energy cost of walking, the calculated whole-body energy cost of walking would increase as well. Hence, the inaccuracies in the model might partly explain why the measured energy cost of walking was, but the calculated energy cost of walking was not higher in older than in young adults. Simulation studies using musculoskeletal models with subject-specific maximal isometric muscle forces and three-dimensional models of muscle contraction are needed to confirm the independent contribution of the walking pattern to the energy cost of walking in young and older adults.

A different walking pattern in young and older adults mediates the influence of Achilles tendon stiffness on the whole-body energy cost of walking. The influence of varying Achilles tendon stiffness on the whole-body energy cost of walking was different in young and older adults when using individualized triceps surae muscle-tendon parameters, and these differences in young and older adults were similar when using generic triceps surae muscle-tendon parameters. This indicates that underlying walking patterns strongly determine the influence of Achilles tendon stiffness on the whole-body energy cost of walking in young and older adults.

In contrast to the effect of walking patterns on the influence of varying Achilles tendon stiffness on the whole-body energy cost of walking, the influence of varying Achilles tendon stiffness on the triceps surae energy cost of walking is strongly affected by lower optimal fiber length in older compared to young adults. The influence of varying Achilles tendon stiffness on the triceps surae energy cost of walking was different in young and older adults when using individualized triceps surae muscle-tendon parameters, but the influence was almost equal in young and older adults when using generic triceps surae muscle-tendon parameters. A single muscle simulation study already reported that an optimal combination of Achilles tendon stiffness and optimal fiber length exists with respect to the energy cost of force production (Lichtwark and Wilson, 2008). This probably indicates that rather the subject-specific combination of triceps surae muscle-tendon parameters than only variations in Achilles tendon stiffness determines the triceps surae energy cost underlying a specific walking pattern.

In young adults, the influence of varying Achilles tendon stiffness on the energy cost of walking was dependent on walking speed. The difference in the influence of varying Achilles tendon stiffness on the energy cost of walking between slow speed and fast or comfortable speed, might be explained by differences in walking patterns. Ankle moments were equal in young and older adults at slow speed, but the ankle moment during push-off was higher in young than in older adults at higher speeds. This indicates that young subjects change their walking pattern with increasing walking speed. The difference in the influence of varying Achilles tendon stiffness on the energy cost of walking between six-minute walk speed and fast or comfortable speed, might be explained by inter-subject variability since the standard deviation is much larger at six-minute walk speed than at other speeds. The six-minute walk speed was subject-specific and it was very high for some subjects (more than 7.5 km/h), but notably lower (below 6 km/h) for other subjects. This variability in walking speeds might cause variability in walking patterns and in the effect of varying Achilles tendon stiffness on the energy cost of walking. This might confirm that changes in walking pattern govern the effect of changes in Achilles tendon stiffness on the energy cost of walking.

Other lower limb muscles or plantarflexor muscles (except the triceps surae muscles) did not determine the influence of Achilles tendon stiffness on the whole-body energy cost of walking in young and older adults. Only when we induced a 50% decrease in Achilles tendon stiffness, the other lower limb muscles’ and plantarflexor muscles’ energy cost clearly increased. Probably, the other plantarflexor muscles compensated for decreased triceps surae force generating capacity due to sub-optimal position on the force-length and force-velocity curves. This was indicated by an increase of the heat production related to shortening lengthening and the heat production related to activation and maintenance with 50% of individualized Achilles tendon stiffness. These energy cost components are strongly determined by the muscles’ normalized fiber length and contraction velocity (Umberger et al., 2003). These increases in heat production were higher than the decreased mechanical triceps surae muscle fiber work, which might be caused by increased Achilles tendon work, when we induced a decrease in Achilles tendon stiffness in older adults. These findings confirm that mechanical muscle and tendon work alone are not sufficient to explain the age-related increase in the energy cost of walking (Monaco and Micera, 2012).

The methods to calculate the energy cost of walking were equally accurate in describing the relationship between whole-body energy consumption and walking speed. This is in agreement with a previous study that also reported small differences in accuracy between different methods to calculate the energy cost of walking only in young adults (Koelewijn et al., 2019). Moreover, in our study, the accuracy of these methods differed greatly between different subjects. Although we analyzed individual correlations between calculated and measured energy cost of walking, we could not consistently attribute inter-subject differences to atypical stride frequencies, body mass, muscle-tendon properties or joint moments during walking. The use of phenomenological Hill-type muscle models might explain why the accuracy of the methods to calculate the energy cost of walking greatly varied between subjects. These phenomenological muscle models do not take into account the physiological process underlying the metabolic cost of active muscle force production, i.e. cross-bridge cycling. Hence, the use of Huxley muscle models that do take into account cross-bridge cycling will probably improve the accuracy of simulation-based methods to calculate the energy cost of muscle force production.

Although the modeling approach allowed us to probe causal relations, it has several limitations. First, Hill-type muscle models are a very simplified representation of the complex muscle-tendon anatomy. For instance, the in vivo Achilles tendon consists of three, partly independent, sub-tendons that arise from the separate gastrocnemius and soleus muscle bellies (Franz et al., 2015). Age-related changes in these sub-tendons’ inter-dependence influence triceps surae muscle-tendon function (Franz and Thelen, 2016). These changes are overlooked in the model that we used in this study because the triceps surae tendons are independent in our models. Moreover, we did only estimate a limited set of triceps surae muscle-tendon parameters. For instance, age-related differences in maximal isometric muscle force were not taken into account (Kemmler et al., 2018). Decreased maximal isometric muscle force requires higher activations for similar force production. Differences in muscle activation are reflected in the muscular energy cost of walking (Umberger et al., 2003). We speculate that individualizing maximal isometric muscle force would increase the plantarflexor and hip extensor muscle activations and energy cost of walking in older but not in young adults. Probably, we would still conclude that walking patterns independently contribute to the energy cost of walking. In contrast, we studied the contribution of Achilles tendon stiffness to the energy cost of walking using within-subject changes in energy cost. Hence, individualizing maximal isometric muscle forces would probably not change the results and conclusions with respect to the independent contribution of Achilles tendon stiffness to the energy cost of walking.

To conclude, Achilles tendon stiffness and different walking patterns in young and older adults independently contribute to the energy cost of walking. Although Achilles tendon stiffness independently influences the energy cost underlying specific walking patterns in young and older adults, its influence is rather limited. Different walking patterns in young and older adults determine the influence of Achilles tendon stiffness on the energy cost of walking in young and older adults. Hence, training interventions should probably rather target specific walking patterns than Achilles tendon stiffness to decrease the energy cost of walking. Based on the results of previous experimental studies, we expected that the calculated hip extensor and whole-body energy cost of walking would be higher in older than in young adults. However, this was not confirmed in our results. Future research might assess the contribution of the walking pattern to the energy cost of walking by individualizing maximal isometric muscle force and by increasing the level of anatomical detail in musculoskeletal models. The interaction between specific walking patterns, Achilles tendon stiffness and the energy cost of walking differs in young and older adults. We suggest that Achilles tendon stiffness minimizes the energy cost underlying older adults’ specific walking patterns. In contrast, it seems that decreasing Achilles tendon stiffness would decrease the energy cost underlying the specific walking patterns in young adults. Furthermore, our results indicate that the age-related differences in triceps surae muscle-tendon parameters determine the age-related differences with respect to the influence of Achilles tendon stiffness on the triceps surae energy cost of walking. Therefore, rather a subject-specific combination of triceps surae muscle-tendon parameters determines the triceps surae energy cost underlying a specific walking pattern.

## Conflicts of interest

None

## Acknowledgments

We thank W. Swinnen, A. Catteau, P. Cortvriendt, B. Scheepers, and T. Wijgaerts for their help during experimental data collection.

